# Mechanism-informed rules tunably balance novelty and feasibility of predicted enzymatic reactions

**DOI:** 10.64898/2026.05.18.726002

**Authors:** Stefan C. Pate, Keith E.J. Tyo, Linda J. Broadbelt

## Abstract

Enzymes catalyze reactions with remarkable specificity and can unlock recalcitrant feedstocks that are dilute, complex, and variable in their constituent molecules. While characterized enzymatic reactions cover a wide range of chemistries, there are an undetermined number of cryptic activities for every known one. These cryptic activities can be elicited through rational design, adaptive laboratory evolution, and increasingly, generative models of proteins. However, prior to tuning a catalyst one must efficiently predict viable novel reactions. In this work we leverage the growing amount of mechanistic enzyme information, specifically the Mechanism and Catalytic Site Atlas, to construct a set of reaction rules that can meet this demand. By explicitly utilizing mechanistic information, the rule sets developed here more accurately identify molecular structures required for catalysis compared to existing curated and heuristically constructed rules. The 899 Distilled rules are constructed directly from characterized mechanisms and cover 62.5% of reactions from Rhea. The Learned rule set is generated from a classifier trained on mechanistic data, allowing full coverage of Rhea and precise identification of mechanism-required atoms (ROC-AUC = 0.98). Additionally, our Learned rules exhibit a more favorable tradeoff between novelty and feasibility and provide users with fine-grained control over this tradeoff. The rules are compatible with all SMARTS-based reaction network expansion and retrosynthesis software.

## Introduction

Enzymes serve an important role in the synthesis of products ranging from commodity chemicals to pharmaceuticals. They exhibit remarkable regio- and stereo- selectivity, function well at mild temperatures and pressures, and, in the cellular context, can accommodate dilute and complex feedstocks. These characteristics have driven their successful utilization in research and industry.^1^ Tens of thousands of reactions and their protein catalysts have been characterized to date,^2,3^ and yet, this may just scratch the surface.

Many undiscovered functions of enzymes are cryptic. These functions, usually referred to as promiscuous, underground, or non-native, tend to be overshadowed by kinetically superior primary activities.^4,5^ However, the slow kinetics of non-native activities are not inevitable; rather they appear to be frozen accidents of evolutionary history. Theories of enzyme evolution suggest that selective pressures drove generalist enzymes toward specificity in many cases, but that enzymes remain labile, which confers evolvability.^5–8^ Leveraging this lability, biotechnologists have modified enzymes to significantly enhance non-native activities, thus engineering systems to synthesize compounds of interest.^9–11^

The underlying enzymatic reaction network inclusive of non-native activities may be very large, making it impractical to store in its entirety. Instead, the edges of the network are represented implicitly by a set of reaction rules, which include only the most important structural molecular features of reactions. In computer-aided synthesis planning (CASP), reaction rules, along with either a target molecule, starting feedstock, or both, are input into a procedure that samples from the underlying reaction network and enumerates linear synthesis paths, trees, or more involved subnetworks.^12–26^ Template-free methods forgo reaction rules in favor of a generative model of reactions^27–31^, but we focus on template-based methods in this study.

The accuracy of reaction rules (or generative model in the case of template-free methods) is critical as incorrect predictions can result in expensive and time-consuming efforts to engineer an enzyme for an activity that was incorrectly predicted, or a desired activity may be abandoned because reaction rules did not correctly predict a viable transformation. An ideal set of reaction rules would, for all possible transformations, capture only the reactant structures necessary for catalysis. Aside from the obvious atoms that have bond changes between reactant(s) and product(s), there are often atoms that temporarily donate or hold an unstable charge in an intermediate step but return to the initial bond configuration in the final product. For example, in the aspartate ammonia lyase reaction, a carbonyl group accommodates a negative charge while a double bond shifts and the amine group leaves, ultimately returning to its initial bond conformation (Fig. 1a). One can then make a very broad set of viable predictions. A minimal description of a reaction is merely the reaction center (RC).^32^ However, in the case of enzymatic reactions, RC-only rules miss adjacent structures which are required for the catalysis and make unreasonable predictions as a consequence. Due to the recursive nature of a network search, this issue compounds catastrophically. To address this issue, previous work has added additional molecular structure surrounding the RC within a topological radius, (RC + R)^33,34^. For example, RC + 1 templates would include all atoms in the RC and bonded to it. In the aspartate ammonia lyase example, the template would include the carbonyl carbons in addition to the RC atoms (Fig. 1a). RC + 2 also includes atoms that are two bonds from the RC, e.g., the entire carboxylic acid groups in aspartate (Fig. 1a). Other approaches use manually-curated reaction rules rather than attempt to extract rules from reaction data.^18,35^ Still other data-drive approaches break with the radial mold in various ways. RDChiral includes in its templates any moieties from a curated list appearing adjacent to the RC.^36–38^ EVODEX differentiates between σ and π bonds as it expands the template about the RC.^39^ The algorithms in EHreact and Ni, 2022 cluster reactions based on common subgraphs, which need not grow radially with respect to the RC.^40,41^

**Figure 1.**
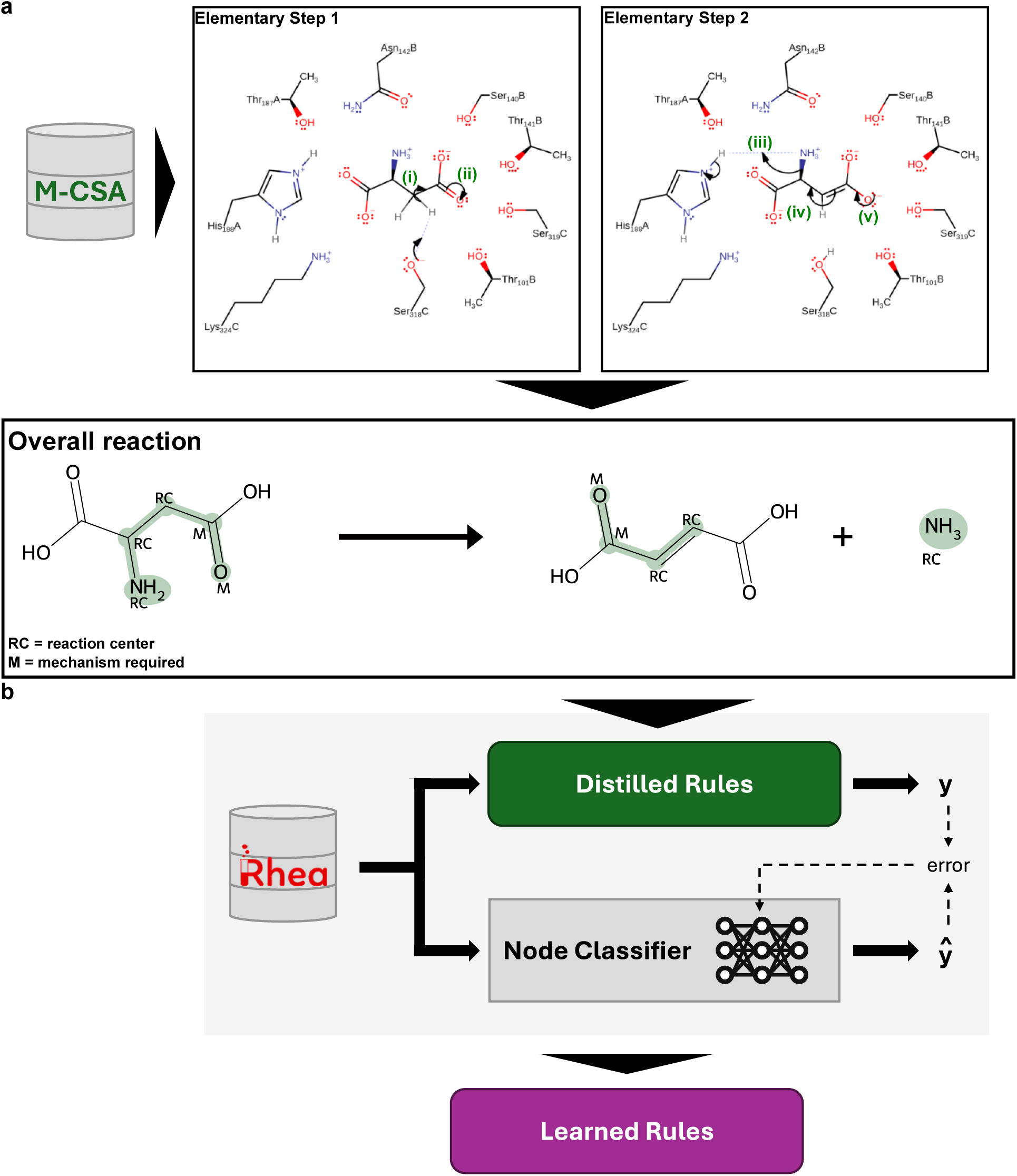
Construction of the Distilled and Learned reaction rule sets. (a) Elementary steps composing overall reactions are downloaded from M-CSA. Curved arrows represent flows of electrons during elementary steps. Some electron flows, such as (i) – (v), involve reactant and product atoms. Some of these atoms comprise the overall reaction center, denoted “RC” in the overall reaction equation. Others are part of only elementary reaction centers but do not undergo transformation between reactant(s) and product(s); these atoms are denoted “M” in the overall reaction. Together, atoms labeled RC and M form the Distilled reaction rule describing the overall reaction highlighted in green. (b) A classifier is trained to label all reactant and product atoms required for the catalysis in Rhea reactions mapped by Distilled rules. Atoms for which the classifier outputs a score above a decision threshold are included in the Learned reaction rules. Abbreviations: RC = reaction center, M = mechanism-required.

Here, we develop a new approach to automatically extract enzymatic reaction rules that directly uses mechanistic data rather than rely on various heuristics to approximate it. Our method was made possible by the high-quality Mechanism and Catalytic Site Atlas (M-CSA), which compiles enzyme reaction mechanisms for a thousand overall reactions^42^. In contrast to other methods which predict elementary steps given an overall enzymatic reaction,^43–45^ we focus on constructing or inferring overall reaction rules based on information from the elementary steps. Our goal is the same as all previous sets of reaction rules, i.e., to capture only reactant structures necessary for catalysis in order to predict viable novel reactions in an efficient manner. We show that our approach more accurately identifies mechanism-required structures and exhibits a more favorable tradeoff between novelty and feasibility of predicted reactions.

## Results

### Curating mechanistic data for enzymatic reaction rule construction and inference

At the time we accessed it, the M-CSA^42^ database contained 1,166 mechanisms for 1,003 overall reactions. 887 of mechanism entries featured detailed arrow-pushing diagrams represented in Chemical Markup Language (CML). The rest were described in natural language. We translated the elementary steps from these detailed mechanisms from CML to SMARTS format, ensuring all reactions were balanced, and consistently atom-mapped throughout all steps. We were able to recover 673 of the 887 CML-encoded mechanisms into balanced, atom-mapped SMARTS. In addition to atom-mapped SMARTS, we store indices of all atoms involved in the mechanism of any elementary reaction step. More precisely, we stored the union of all elementary reaction centers. This is a superset of the overall reaction center and comprises a template of a valid reaction rule (Fig. 1a). We then reversed all 673 balanced, atom-mapped SMARTS leaving 1,346 reactions with all reactant, product, and active site residue structures specified. Finally, we extract just the structures involved in the mechanism to form rule templates and eliminate duplicate rule templates. We refer to the resulting collection of 899 unique rules generated through this procedure as the Distilled rules.

With the Distilled rules in hand, we next tested their coverage of known biochemistry as described in the Rhea database^2^. The Distilled rules recapitulate 62.5% of atom-mapped enzymatic reactions from the Rhea database^2^ (Table 1). The vast majority of these recapitulated reactions (95%) are *not* found in the M-CSA database, making them useful for reaction prediction and retrosynthesis tasks on their own. To cover the remaining 37.5% of reactions not mapped by the Distilled rules, we leveraged the mapping of Distilled rules to the overall enzymatic reactions they correctly recapitulated to construct a labeled dataset. We then used this rules-to-reactions mapping to train a parametrized model that can be used to create reaction rules for enzymatic activities not in the M-CSA database (Fig. 1b). The model approximates the mapping of the Distilled rules but is not limited to the 899 templates of this rule set. It accepts any overall enzymatic reaction, and outputs what can be formatted into a reaction rule, achieving full coverage of all atom-mapped overall reactions. We will build up to this rule set, referred to as the Learned rules, in subsequent sections.

**Table 1.**
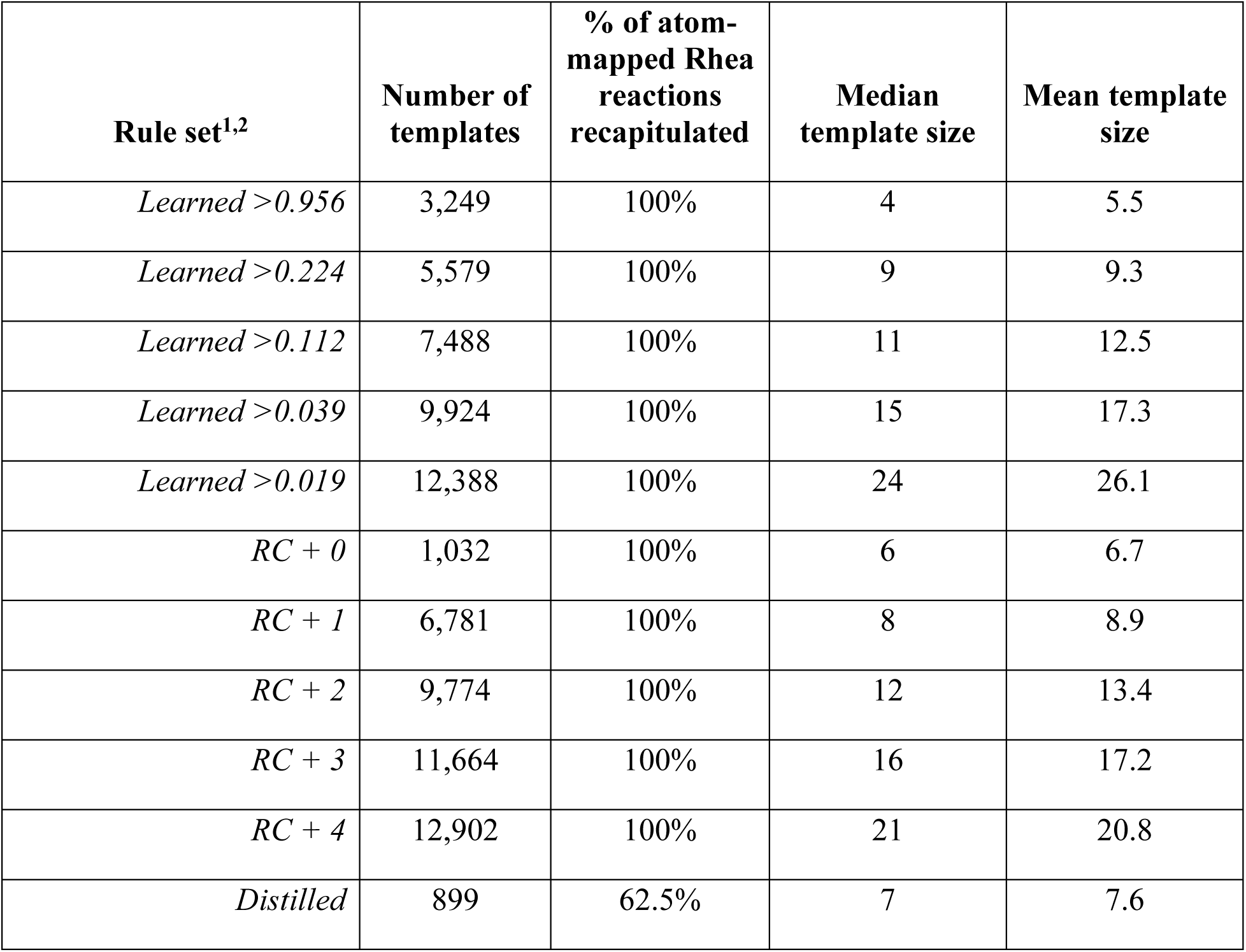
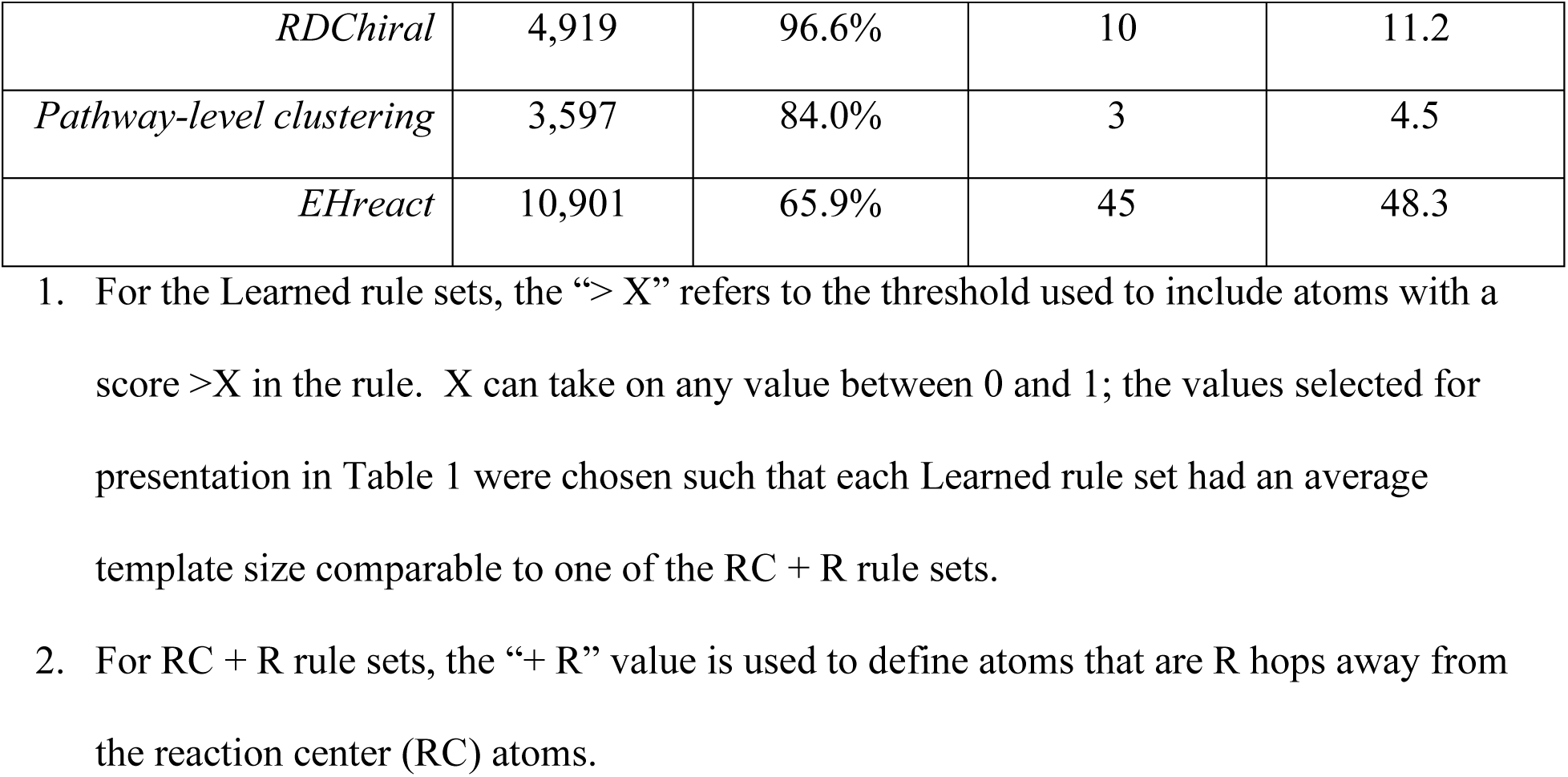
Summary statistics for evaluated rule sets.

### Mechanism-required atoms are sparsely distributed over multiple distances from the reaction center

An analysis of enzyme reaction mechanisms reveals the limitations of using an RC + R approach to constructing reaction rules and instead motivates an approach that utilizes mechanistic information. While it is true that mechanism-required atoms tend to be within a few bonds of the reaction center, with nearly 90% located within a single bond (Fig. S5a), it is not sufficient for a rule extraction method to include merely the majority of required atoms, aggregately across many reactions. An ideal method would include all atoms required for the mechanism per each reaction while minimizing the amount of unnecessary spectator atoms included in the rule template. This goal is made challenging by the fact that the distance from the reaction center of the most distal of the mechanism-required atoms varies reaction to reaction. The maximum distance varies between zero and four for 98% of all reactions with described mechanisms (Fig. 2a). While 90% of mechanism-required atoms are located within a single bond of the overall reaction center, RC + 1 rules would only fully cover the required structural elements for 63% of all reactions (Fig. 2a). At the same time, 81% of atoms in RC + 1 are unnecessary for the mechanism (Fig. 2b). The tradeoff between covering all required atoms and rejecting spectator atoms, i.e., those that are not involved in the mechanism, is harsh for the RC + R strategy. With an RC + 4 rule set, all mechanism-required atoms for 98% of characterized reactions are included, but at the cost of including 3.9 spurious atoms for every required one (Fig. S5b). These overly prescriptive reaction rules do not generalize much beyond known enzymatic reactions. However, relative to the Distilled rules derived from M-CSA, which only covered 62.5% of known reactions, RC + 4 rules have the advantage of very high reaction coverage for known reactions. We sought a data-driven solution to this issue that would both have high coverage of known reactions while minimizing the spurious atoms to allow a higher degree of generalization to suggest novel reactions.

**Figure 2.**
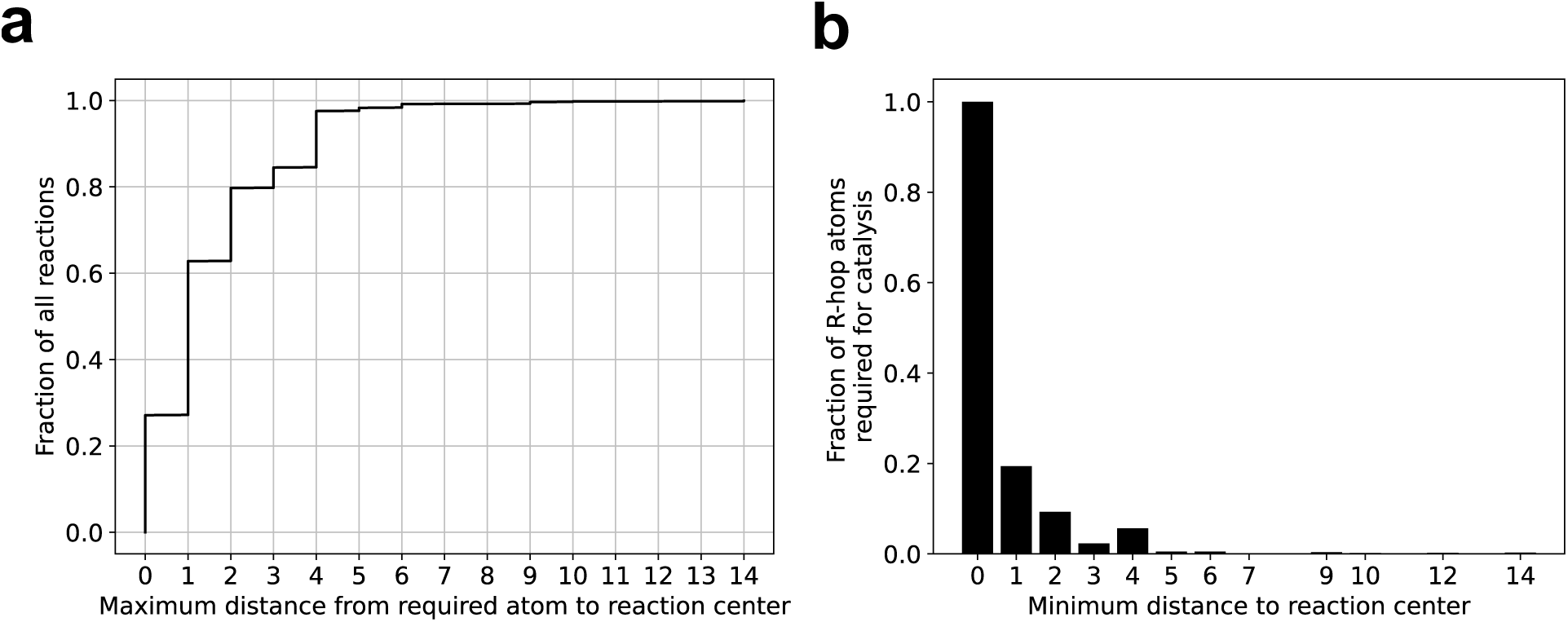
Mechanism-required atoms are sparsely distributed across multiple topological distances from the RC. (a) A CDF of fraction of overall reactions in the M-CSA database against the maximum distance between overall reaction center atoms and mechanism-required atoms. (b) A plot of the fraction of atoms required for catalysis out of all atoms of a given minimum topological distance (# of bonds) from the overall reaction center, *R*. Fractions at *R > 6* are all less than 0.003.

### Learning a parametrized approximation of mechanistically relevant structural patterns

Ultimately, we desired a set of enzymatic reaction rules informed by mechanistic information which generalize to as many known overall enzymatic reactions as possible, and with an added requirement of granting user control over level of rule specificity. We achieved this by first leveraging the Distilled rules to create a labeled dataset and then formulating reaction rule writing as a binary classification task that we solved with machine learning.

The number of enzyme reaction mechanisms retrieved from M-CSA was relatively small. Nevertheless, through mapping the derived Distilled rules to Rhea, we assembled a dataset of 12,137 reactions with atoms labeled as required for the mechanism or not. When a reaction rule is mapped to a full reaction, in addition to recapitulating the reaction (transforming the reactants into the observed products), one also obtains a correspondence between the rule’s template and the reactants’ molecular structures. The resulting data labels are binary, and there is one label for each atom in the full overall reaction. For an example, see Figure 3 in which atoms of the positive class – involved in the mechanism – are highlighted green.

**Figure 3.**
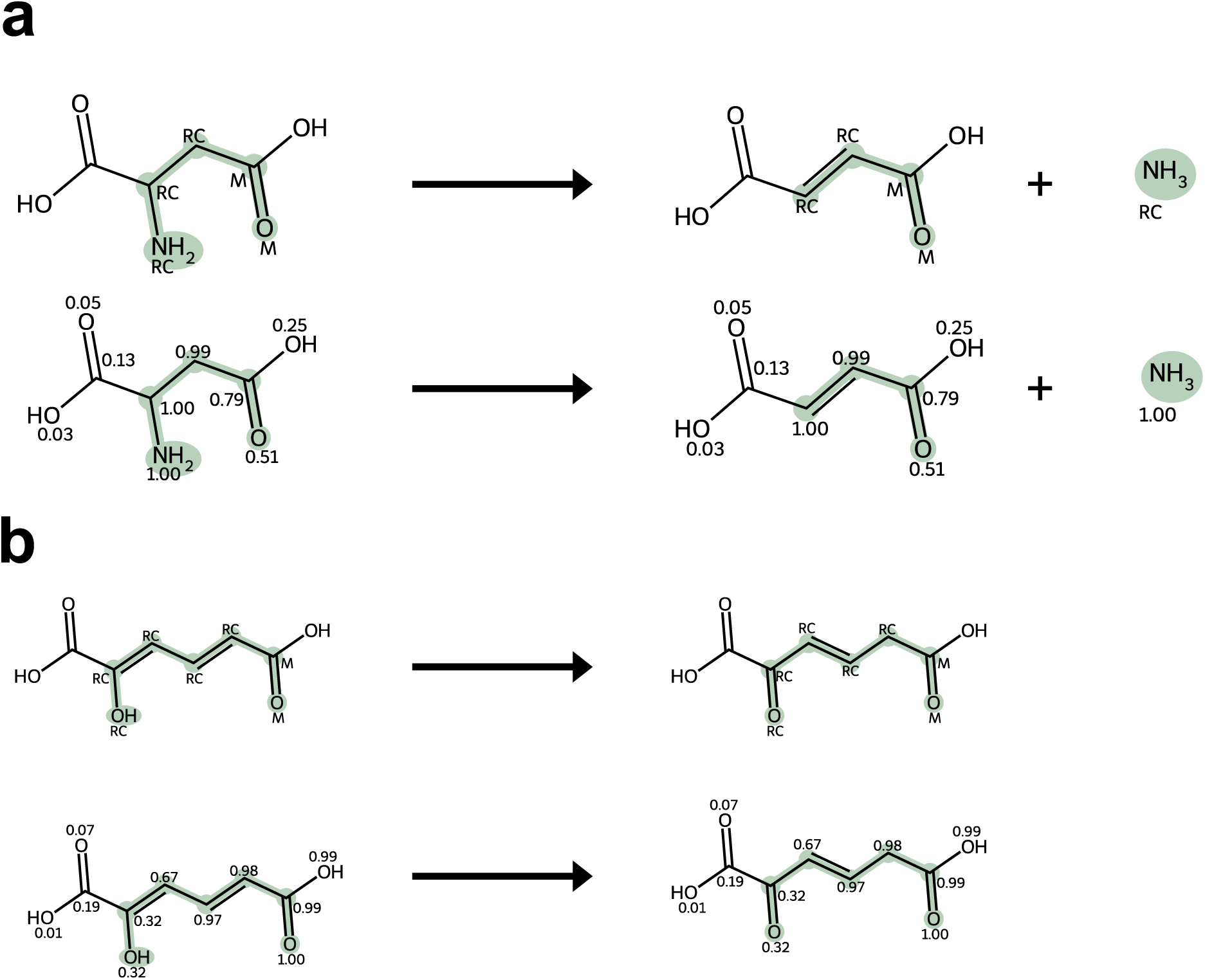
Model-inferred labels closely adhere to Distilled templates in difficult examples. (a) An example of a highly successful labeling of a difficult overall reaction from M-CSA. Top, aspartate ammonia lyase reaction (EC 4.3.1.1) highlighting atoms involved in any elementary step of the catalysis. Bottom, the same reaction visualizing the scores output by a trained model. This model was not trained on any reaction mapped by the Distilled template displayed in green. Note that the scores are the same for a given atom in both the reactants and products, as the score is specific to the atom. (b) An example of a reasonably successful labeling of a difficult overall reaction from M-CSA. Same format as (a) but with 2-hydroxymuconate tautomerase reaction (EC 5.3.2.6). Abbreviations: RC = reaction center, M = mechanism-required.

Given the structure of this dataset, we constructed a model that accepts a featurized reaction graph as an input, and outputs as many binary labels as there are atoms in the reaction (Fig. 3).

The classification task associated with this form of dataset can be viewed in multiple lights. One can think of a reaction as a data point, making the task a form of multilabel binary classification in which the number of labels, equal to the number of atoms, is variable. Alternatively, one can think of an atom as a data point, and data points are batched together depending on molecular structure. A third interpretation of the task as node classification emphasizes the graphical nature of the data. Motivated by this, we employed GNNs to learn vector representations of the atom nodes, dependent on their structural surroundings, and fed these vectors into a FFN to output scalar scores that were later made discrete labels by application of a threshold.

We evaluated the model’s generalization to mechanisms not present in the M-CSA dataset through the use of group 5-fold cross validation in which reactions were first grouped by the mapped Distilled rule prior to being split into the test dataset. We employed group 3-fold cross validation nested within the 5 outer splits for hyperparameter optimization. See Methods for details.

We were most interested in decision-threshold-invariant metrics as we designed to grant users control over the value of the decision threshold to tune specificity to their needs. Considering first the interpretation of atoms as data points, the trained node classifier achieves a more favorable tradeoff between precision and recall than the RC + R strategy where R is the specificity control parameter for this scheme (Fig. 4a). The area under the precision-recall curve (PRC-AUC) for the former was 0.95 while that of the latter was 0.87. On this task, RDChiral^36,37^ performed similarly to RC + 1. The interpretation of reactions as data points suggests a slightly different evaluation in which successful recall of a data point means all mechanism-required atoms for a given reaction are labeled with the positive class. The definition of the precision metric is unchanged. Here again, the trained node classifier outperforms heuristic algorithms with PRC-AUC of 0.85 vs 0.59 for the RC + R method (Fig. 4b). Similar results were observed when the node classifier was trained and evaluated exclusively on reactions found in the M-CSA dataset. In this case, the node classifier achieved an atom-wise PRC-AUC of 0.93 vs RC + R with 0.89 and a reaction-wise PRC-AUC of 0.76 vs RC + R with 0.64 (Fig. S9).

**Figure 4.**
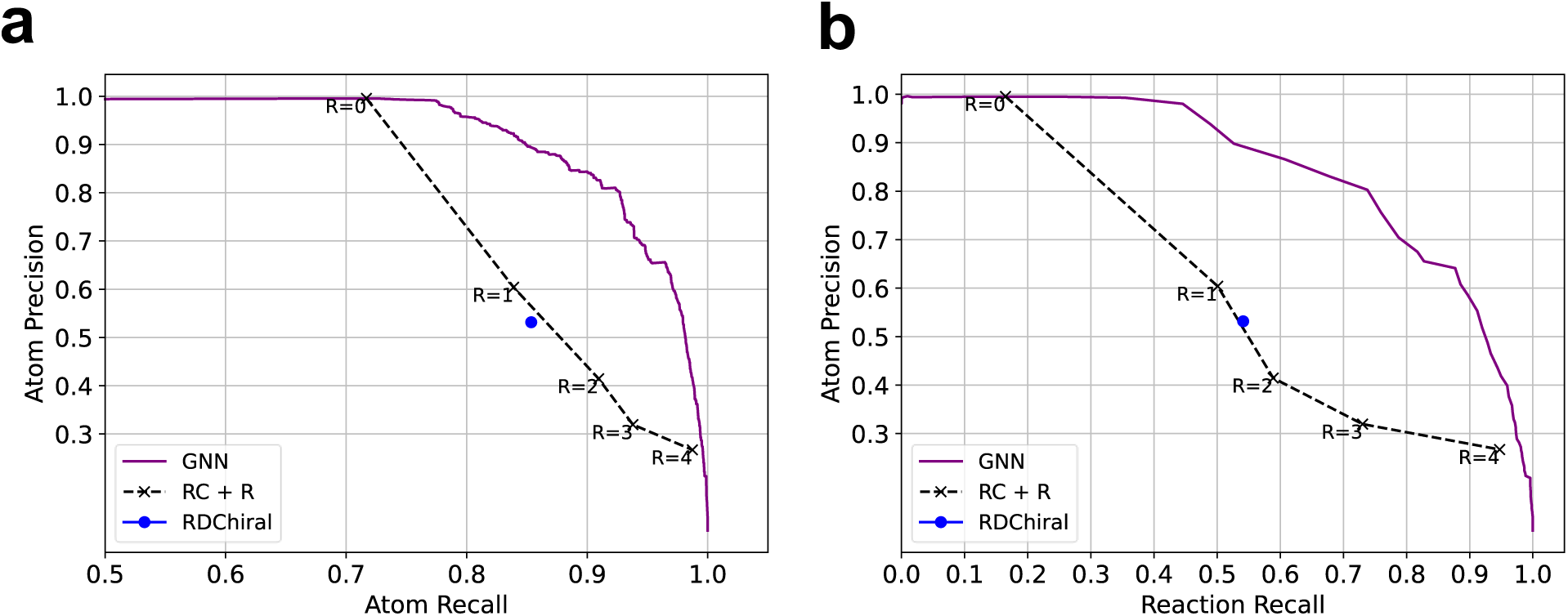
GNN model successfully labels atoms putatively required for the catalysis of enzymatic reactions. (a) Precision-recall curves for the task of labeling atoms as required for the catalysis of an overall reaction. (b) Precision score evaluated for labeling atoms required for the catalysis versus recall evaluated for full overall reactions. For the latter metric, a correctly recalled data point is defined as an overall reaction with all atoms required for the catalysis of that reaction correctly labeled as with the positive class.

Two examples illustrate how the trained node classifier outperforms RC + R on the binary classification of mechanism-required atoms (Fig. 3). Both reactions shown in Fig. 3 are catalyzed via mechanisms that involve atoms outside of the overall reaction center, so reaction center only rules (RC + 0) miss this critical information. Even increasing specificity to RC + 1 leaves out required carbonyl oxygen atoms in both examples and at the same time includes unnecessary backbone carbons in the template. Increasing specificity further to RC + 2 is even more problematic as now all atoms in the reactants are required by the resulting template, thus precluding the prediction of any novel reactions which are not a superset of the observed structure.

In contrast, it is possible to generate more tailored reaction rule templates by tuning the decision threshold of the trained node classifier to assign the atoms to classes. In the best cases, scores given to mechanism-required atoms are much higher than those given to spectator atoms (Fig. 3a). Even in cases where the model makes an error by giving a high score to an unnecessary atom, it fails more gracefully than the RC + R method (Fig. 3b).

### Generating Learned rules with a trained node classifier

The successful training and evaluation of the node classifier on Distilled-labeled data lay the groundwork for the Learned rule set(s). The procedure consists of labeling atoms from overall enzymatic reactions, extracting structural patterns corresponding to these labeled atoms in SMARTS, and enumerating all unique rule templates. As a demonstration of this procedure, we generated variants of the rule sets for a range of decision thresholds, which is inversely proportional to the resultant rules’ specificity. The number of unique templates in the rule set and the average template size increase concomitantly with specificity (Table 1). Beyond spanning a range, we selected decision thresholds to approximately match template size to the five variants of RC + R rules evaluated in this work (Fig. 5a). We provide evidence of the superior coverage of known enzymatic reactions by the Learned rules, relative to Distilled, in the form of an aggregate count of reactions recapitulated, which equals that of the reaction center only rules (RC + 0) (Table 1), as well as broken down by Enzyme Commission (EC) number as a proxy for the diversity of chemical transformations covered. We provide this breakdown up to the first EC digit (Fig. 5b) and second EC digit (Fig. S4) for a more granular view.

**Figure 5.**
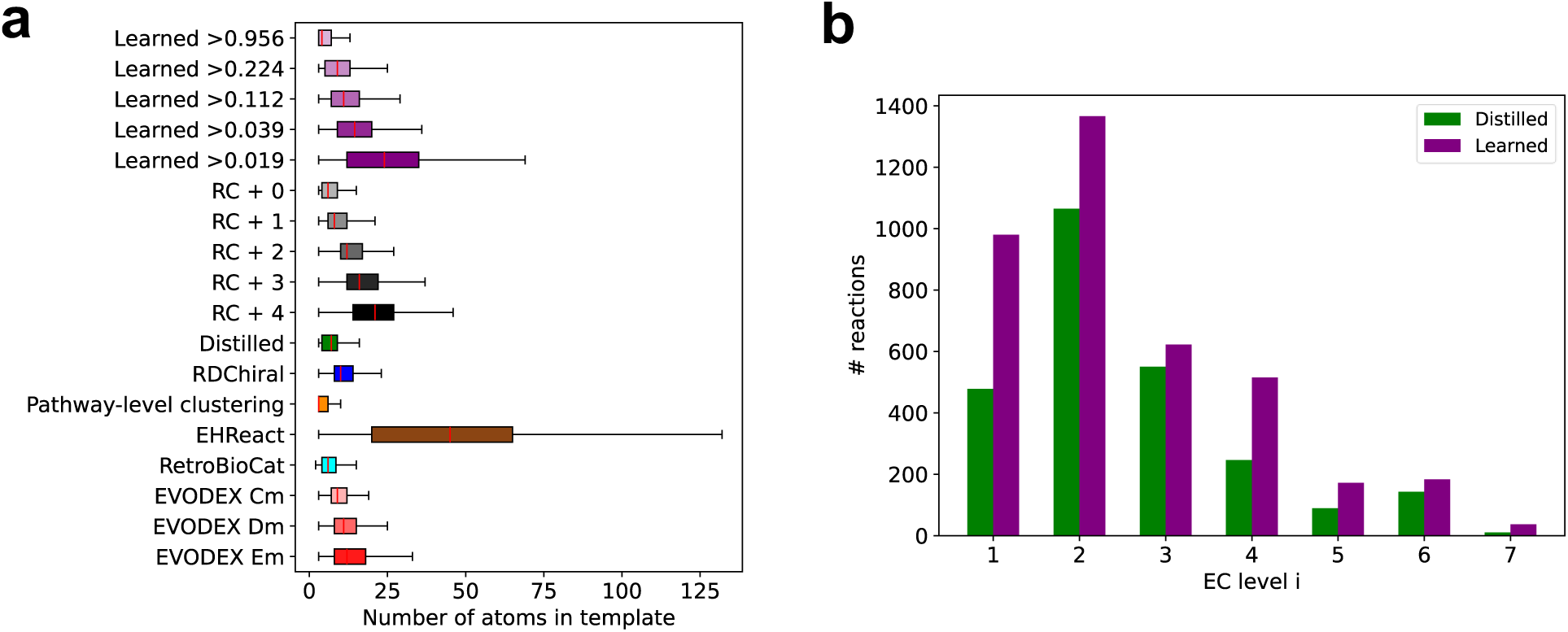
Summary of the specificity and coverage of the Distilled and Learned rule sets. (a) Box and whisker plots depicting the distribution of template sizes, defined as the number of atoms included, from each set of reaction rules that were evaluated in the test reaction network expansions summarized in Figure 6. Median template sizes are represented by red lines and are the same values presented in Table 1. Other ticks represent (from left to right) minimum, first quartile, third quartile, and maximum. (b) Comparison of the coverage of the Enzyme Commission (EC) number classes (i.j.k.l) by the Distilled and Learned rule sets. The y-axis counts the number of unique reactions mapped which are assigned to the EC i digit plotted on the x-axis. Note that all Learned rule sets cover the same known reactions.

### Comparison of mechanism-informed and established rules in reaction prediction

Prediction of novel enzymatic reactions is one of the main applications of enzymatic reaction rules. Often this is performed iteratively to compose reactions into multi-step synthesis pathways. The ideal set of reaction rules would predict many novel reactions that extend the current reach of enzymatic synthesis while minimizing the number of infeasible reactions predicted which strain processing and memory resources without delivering results that can be realized experimentally. These costs compound with each subsequent reaction network generation step which multiplies the size of the network. To evaluate rule sets’ ability to efficiently predict feasible novel reactions, we ran two test network expansions.

In the first network expansion, we seeded the network with a set of starting molecules designated by the Agile BioFoundry as “beachhead molecules” in a graphic adapted from Lee, S.Y., et al., 2019 (Fig. 6a). We then applied reaction rules for two generations to the starting molecules and the progeny in the first generation to create a growing reaction network. For each rule set and corresponding generated reaction network, we counted the number of novel predicted reactions and the fraction of those predictions labeled as feasible by an existing discriminative model, DORA-XGB^46^. We did not conduct this analysis on RetroBioCat or EVODEX rules since they generate unbalanced reactions and DORA-XGB was trained exclusively on balanced reactions.

**Figure 6.**
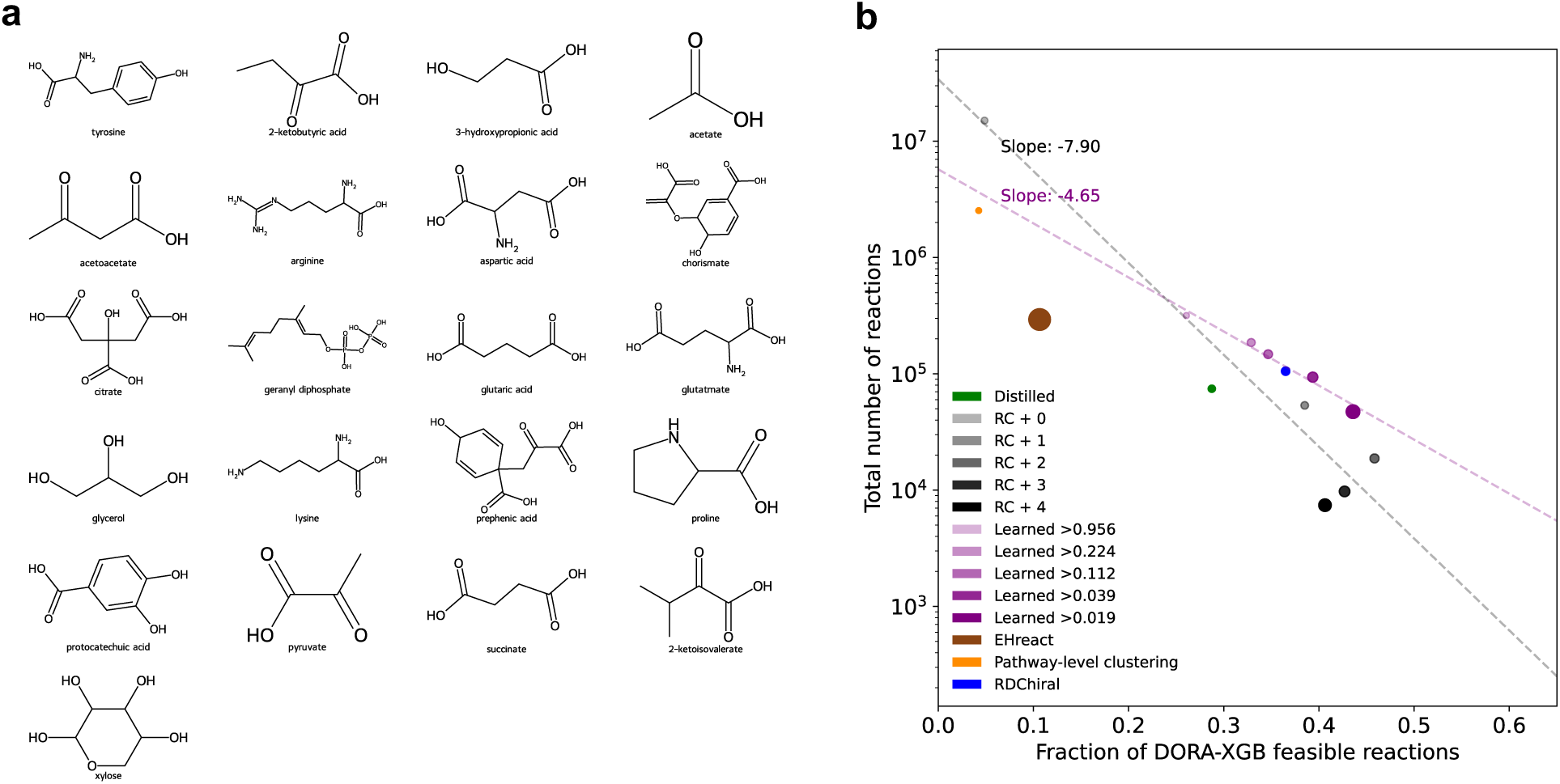
Learned rules exhibit more favorable novelty-feasibility tradeoff than RC + R. (a) The molecular structures of the 21 starting molecules which seeded the test network expansion^1^. (b) Total number of unique predicted reactions plotted against the fraction of total reactions labeled as feasible by the DORA-XGB classifier^46^. Each point corresponds to one reaction network expansion and corresponding reaction rule set used for the expansion. Point size scales linearly with the median template size of the reaction rule set. Dashed lines represent lines of best fit for the RC + R points and Learned points.

An exponential function fit well the novelty-feasibility tradeoff for both rule sets with a tunable parameter controlling specificity, Learned with R^2^ = 0.98 and RC + R with R^2^ = 0.94 (Fig. 6b). The decay rate for the Learned rule sets, −4.65, quantifies a more favorable tradeoff relative to the RC + R rules whose decay rate was −7.90. The Distilled rule set achieves feasibility between the most and second most permissive Learned rules but with significantly fewer reactions predicted (Fig. 6b). This may be attributed to either the relatively low coverage of enzymatic transformations of the Distilled rule set, particular biases of the discriminative feasibility model, or both. The RDChiral rule set resulted in a reaction network very close to the Learned novelty-feasibility tradeoff line, indicating that the manually curated set of important functional groups with which it augments the reaction center is well aligned with mechanistically relevant structural patterns (Fig. 6b). Finally, a striking difference between the Learned and RC + R rule sets was the sensitivity to their respective specificity control parameters. For example, increasing from RC + 0 to RC + 1 dramatically reduces the number of reactions 281-fold and increases the fraction which are feasible 8-fold (Fig. 6b). By comparison, tuning the decision threshold of the Learned rules yields much finer-grained control over specificity.

In the second test network expansion, we sought a method that does not rely on a probabilistic model to score reaction feasibility. We achieved this with a time-based split wherein we constructed rule sets based on all reactions first characterized before a cutoff date. Then we performed a one-step expansion, seeded with all compounds involved in reactions characterized after this cutoff date. For each rule set, we quantified the fraction of post-cutoff reactions predicted by pre-cutoff rules and the ratio of post-cutoff characterized reactions to total reactions predicted. In total, 4,466 reactions were characterized after a cutoff date of 2015 (Fig. 7a). As not all reaction centers (alternatively RC + 0 rules) had been characterized by 2015, even the RC + 0 rules could not predict all post-2015 reactions. These rules predicted 60.2% of post-2015 reactions and did so inefficiently; just 0.035% of all reactions generated were experimentally characterized post-2015 (Fig. 7b). The most permissive of the Learned rules (>0.932) predicted 56.4% of post-2015 reactions and were ten times more efficient than the RC + 0 rules (Fig. 7b). In general, the Learned rules exhibited a more favorable tradeoff between coverage of empirically characterized rules and efficiency than the RC + R rules. The RDChiral rule set performed comparably to the second most permissive Learned rules (>0.069). Most of the EVODEX variants performed comparably to the RC + 1 rules. The RetroBioCat and EHreact predicted significantly fewer post-2015 reactions than the other rule sets tested.

**Figure 7.**
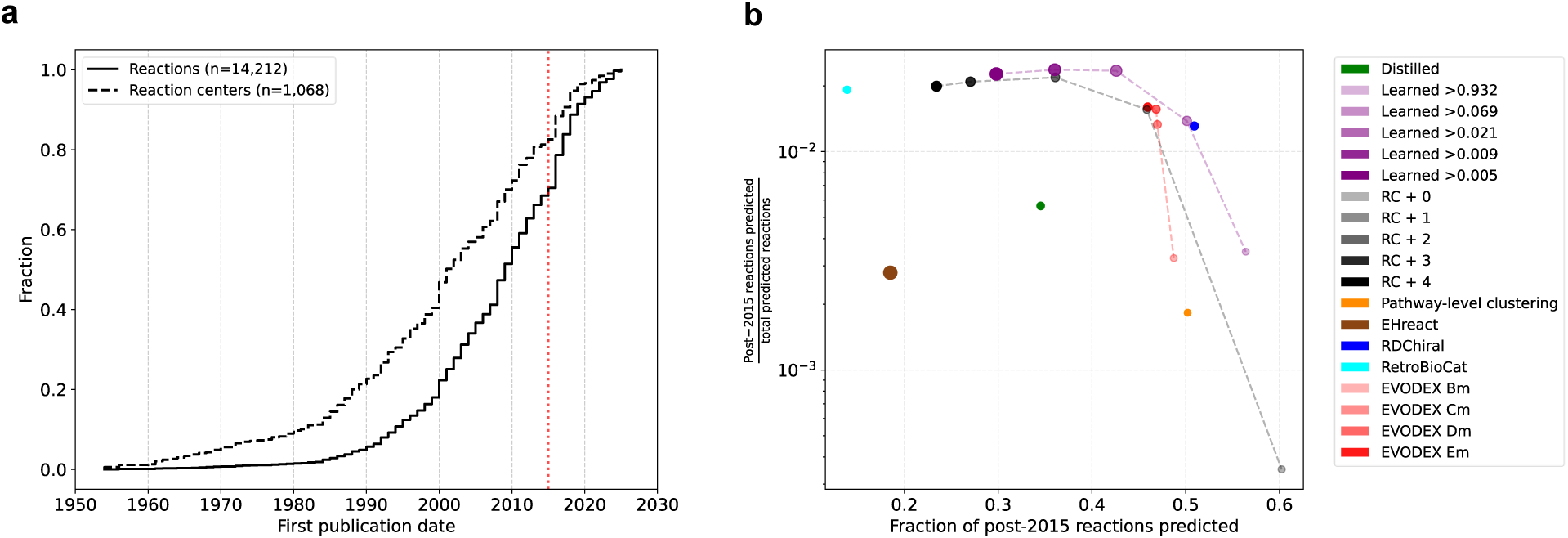
Learned rules effectively balance coverage and efficiency in a time-based split. (a) CDF showing first publication date for unique data set reactions and reaction centers (equivalently RC + 0 rules). The cutoff date chosen for the time split, 2015, is shown in red. (b) Tradeoff between fraction of post-2015 characterized reactions predicted by pre-2015 rule sets and the ratio of post-2015 characterized reactions to all reactions predicted by each rule set. Dashed lines connecting the Learned and RC + R rule sets serve as a visual aid of the achievable tradeoffs of each set.

The reactions predicted by the RC + 0 rules are a superset of reactions predicted by all the other rule sets, but using this rule set comes at a severe cost due to the sheer volume of reactions its rules predict. Thus, compared to the most expressive of the practical RC + R rules, RC + 1, we highlight that both the Distilled and Learned rules accessed a distinct set of predicted reactions (Fig. 8a). We provided two examples of such predicted reactions. In the first, the Learned rules predict a reaction whose nearest known analogue is a 4-hydroxy-2-oxovalerate aldolase reaction (EC 4.1.3.39) (Fig. 8b). The RC + 1 rules extract a template based on this known analogue that specifies a primary carbon alpha to the aldehyde moiety, so it does not generalize to the predicted reaction which has an alpha ketone group. The second example, predicted by the Distilled rules, shows a dehydration reaction in analogy to an arogenate dehydratase reaction (EC 4.2.1.91) (Fig. 8c). Here again, the RC + 1 rule includes a secondary carbon one bond from the reaction center which is unnecessary for the mechanism and fails to generalize to the predicted reaction.

**Figure 8.**
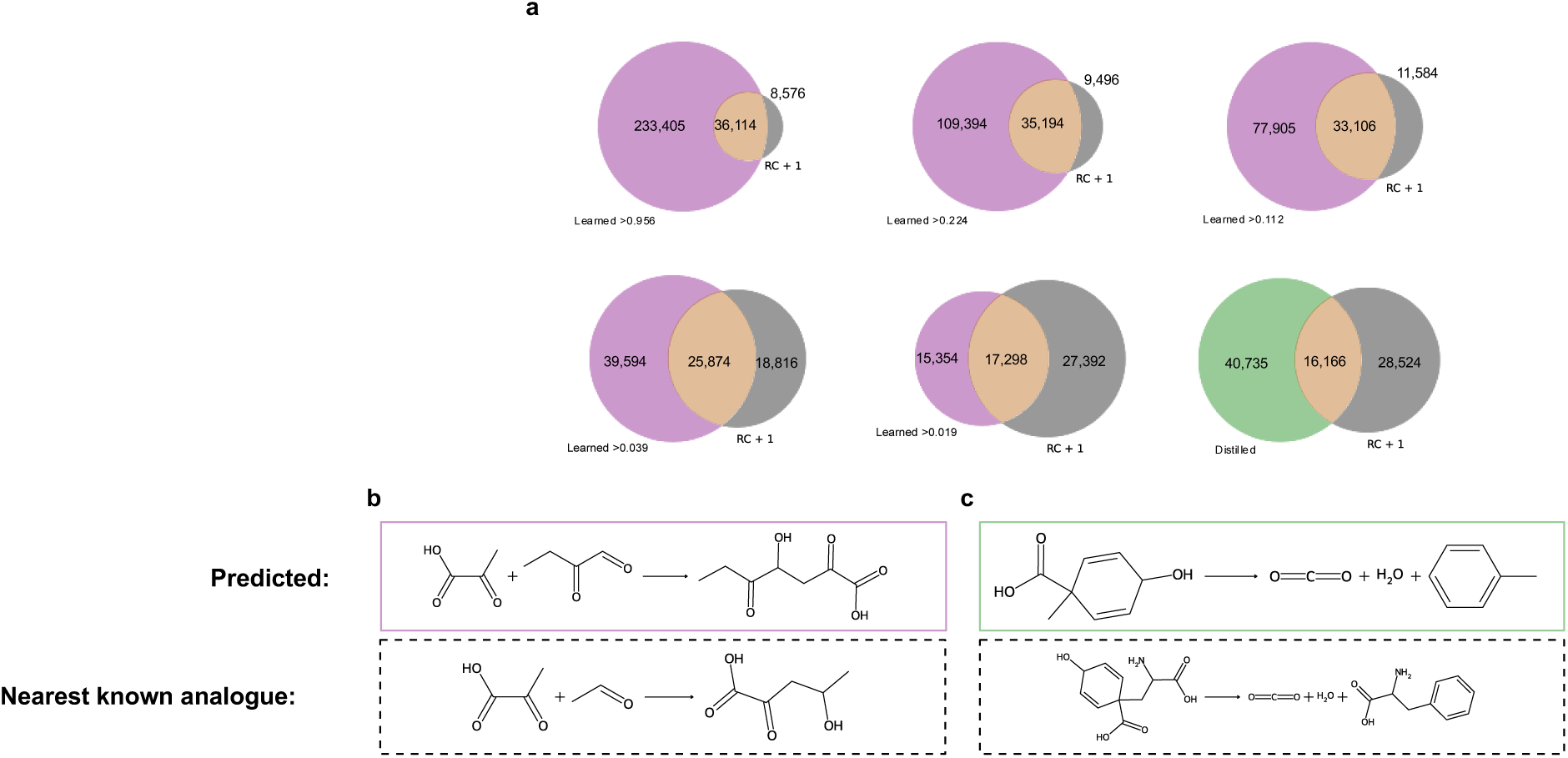
Learned and Distilled rules predict distinct sets of reactions from RC + 1. (a) Venn diagrams contrasting the predicted reactions of the Learned rules or Distilled rules with those predicted by the RC + 1 rules. Purple = Learned, Gray = RC + 1, Tan = Intersection. (b) Top, an example reaction predicted by a rule from the Learned >0.019 set. Bottom, the nearest known analogue, 4-hydroxy-2-oxovalerate aldolase reaction (EC 4.1.3.39), of the predicted reaction. (c) Top, an example reaction predicted by a rule from the Distilled set. Bottom, the nearest known analogue, arogenate dehydratase reaction (EC 4.2.1.91), of the predicted reaction.

## Discussion

In this work, we developed two sets of reaction rules which are novel in their use of mechanistic data. We demonstrated, relative to common heuristics, a trained node classifier more accurately identifies mechanism-required molecular structures, extending the coverage of mechanism-informed rules beyond reactions with characterized mechanisms. Additionally, the Learned rules exhibit a more favorable tradeoff between novelty and feasibility of predictions, evidenced by a less negative exponential decay rate. By varying the decision threshold on the node classifier, users can tune rule specificity with more fine-grained control than by varying the radius in an RC + R scheme. Furthermore, our mechanism-informed workflow stands to improve with additional characterization of enzyme reaction mechanisms, unlike prior methods.

In the broader context of computer-aided enzymatic synthesis planning, our rules are designed to be interoperable with most network expansion and retrosynthesis tools as they are encoded in the widely used SMARTS format. Building the correct information into reaction rules (or generative model) will always be more efficient than methods which predict and then filter with a discriminative model or heuristic. Our method shares many of the limitations of other reaction rules and template-free reaction predictors. It does not account for noncovalent interactions between reactants and amino acid residues in the enzyme active site which will impact activation energies and thus kinetics through processes such as binding and transition state stabilization. These interactions heavily depend on the protein, which one can in various ways engineer to accommodate the required transformation. Nor does our method take thermodynamics into account, which depends on reactant and product concentrations in addition to their structures. However, by increasing the quality of reaction predictions, our method saves computational resources for these important downstream considerations, and moreover minimizes unnecessary experimental effort, thereby making enzymatic synthesis planning more efficient overall.

## Methods

### Curating and processing data

Overall enzymatic reaction data were downloaded from Rhea.^2^ To standardize the reaction SMILES,^47^ we removed stereochemistry, neutralized charges that vary depending on environmental conditions (e.g., protonating -OH groups), applied standard molecule normalization transformations, and canonicalized molecular SMILES. Standardization was carried out with cheminformatics library RDKit.^48^ All transport reactions (e.g., between cellular components) were removed. We automatically generated RCs for each reaction by applying a set of minimal enzymatic reaction operators^32^ to reactants, alternately protecting all but one substructure matching the operator’s RC template.

We removed stereochemistry to enable the use of the DORA-XGB^46^ tool to compute feasibility scores. However, it is not necessary to do so; our template extraction process supports stereochemistry. As a demonstration of this, we extracted chiral templates from five reactions with stereochemical information and successfully recapitulated the reactions using RDChiral’s rdchiralRunText function^36^ (Fig. S8).

Mechanistic enzymatic reaction data were downloaded from the Mechanistic & Catalytic Site Atlas (M-CSA).^42^ Elementary reaction steps and arrow-pushing diagrams were converted from Chemical Markup Language (CML) to SMILES using a custom script. We extracted reactants and products from each step, discarding all amino acid residues and cofactors. We then ensured that each elementary step was fully balanced and consistently atom-mapped. Finally, we generated atom-mapped SMILES for the overall reaction consisting of the elementary steps, and stored indices of atoms involved in any elementary step, including primarily those adjacent to bonds which changed order in the step, i.e., the elementary step’s reaction center, and atoms involved in coordinate bonds, e.g., with metal ions. These atom indices were used subsequently in the creation of Distilled reaction rules and the construction of the binary classification task ultimately leading to the creation of the Learned reaction rules.

### Assembling reaction rules

Reaction rules were encoded as SMARTS. The following procedure was followed for RC + R rules, Distilled rules, and Learned rules. For atoms included in reaction rule templates, we specified features: degree, valence, atomic number, formal charge, membership in ring structures, membership in aromatic structures, and number of heteroatom neighbors. Additionally, in the case where reaction rule templates contained disjoint subgraphs within single molecule templates, we connected all subgraphs to the reaction center via anonymous atoms. Anonymous atoms do not have any specified features and thus can match any atom. Bonds in which an anonymous atom was involved were also anonymous; in other words, no features of these bonds were specified. Atom and bond anonymity was achieved with wildcard characters “*” and “∼”, respectively. We specified the bond order of bonds occurring between two non-anonymous atoms.

The criteria for inclusion of an atom non-anonymously in a reaction rule template is as follows for different classes of reaction rules. Distilled: all atoms whose indices were saved during the processing of M-CSA data, i.e., part of any elementary step reaction center or coordinate bond. Learned: all atoms labeled with a score above the chosen decision threshold by a trained GNN classifier described further in a section, “Binary classification of atom involvement in catalytic mechanism”. RC + R: all atoms located within R bond hops of the reaction center. RDChiral^36^ and EVODEX^39^ reaction rules were extracted from the portion of Rhea overall enzymatic reaction for which we had atom mappings. We applied the template extraction pipelines provided in the publicly available code accompanying each of those publications. Pathway-level clustered reaction rules were constructed by first clustering reactions with similar reactant structures and then writing a reaction rule which includes the maximum common subgraph of all reactions grouped into a cluster. For more details see Ni, 2022.^41^ EHreact^40^ rules were generated by applying their pipeline to the Rhea reactions we curated without all-atom mapping. We used the code accompanying the EHreact publication, based on Reaction Decoder Tool^49^, to get all atom mappings. To generate a final set of rules, we selected the most specific mutual template of each tree as recommended in EHreact.

Finally, we ensured that all reaction rules were unique.

### Mapping rules to reactions

Using the rdChemReactions module of RDKit, we attempted to apply each reaction rule to each overall reaction from Rhea. If the rule was able to recapitulate the reaction, i.e., generated the observed product(s) from the provided reactant(s), that rule was said to “map” that reaction. In the case of multiple rules mapped to a single reaction, we resolved the conflict with one of two approaches. For the RC + 0 rules, we preferred rules without structural patterns consisting of disjoint fragments of a single whole molecule, and we preferred rules without C-C bond-breaking. After applying these criteria, if there were still multiple mappings, we chose the rule which mapped more overall reactions. For the case of all other reaction rule sets, we selected the rule with the largest template, in terms of number of atoms included in that template.

We modified the rule-reaction-mapping criteria in the case of RetroBioCat and EVODEX rules, which are not always atom-balanced. For these rule sets, we aggregated all generated reactions which mapped subsets of known reaction’s reactants to subsets of its products. In other words, we took the union of generated reactions’ reactants and the union of generated reactions’ products. We then checked that every reactant and product from the known reaction appeared in the left-hand side and right-hand side unions, respectively. If this condition was met, the known reaction was counted as “mapped”. Furthermore, we allowed for common currency molecules, such as oxygen, water, or the pair NAD^+^/NADH to be omitted.

### Binary classification of atom involvement in catalytic mechanism

#### Data labeling

We first mapped overall enzymatic reactions from Rhea with the assembled Distilled rules. For this subset of mapped reactions, we labeled each atom included non-anonymously in the Distilled rule with the positive class, “involved in the catalytic mechanism”. All other atoms in the reaction were labeled with the negative class.

#### Model construction

We constructed GNN-based classifiers using Chemprop^50^ and PyTorch Lightning. We used the default atom and bond features of Chemprop and chose the condensed graph of reaction^51^ featurizer.^52^ Input features were transformed into hidden representations via bond-wise message passing layers^53^ and subsequently fed into a custom prediction head (either linear or feedforward neural network) which output scalar values for each atom in the input reaction.

#### Training

All model parameters were learned using a binary cross entropy loss function with a positive weight parameter chosen through hyperparameter optimization.

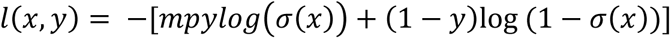

Where m = negative multiple, p = positive multiplier, y = label, x = model output. We used an Adam optimizer and Noam learning rate schedule with linear warmup.^50,54^

#### Hyperparameter optimization (HPO) and evaluation

We conducted HPO and subsequently evaluated models using nested cross validation with three inner folds and five outer folds. Data were split using the GroupKFold class from scikit-learn’s model_selection module.^55^ Reaction data points were grouped by which Distilled reaction rule mapped them. We performed Bayesian HPO using the package Optuna with which we maximized the area under the precision recall curve.^56^ All optimized hyperparameters and the corresponding values used in the evaluated and production models are provided in Table S1.

#### Labeling atoms as input to Learned rules

Using the trained production models, we predicted labels for atoms in all overall reactions from Rhea for which we had atom mappings, which is required for featurization as a condensed graph of reactions. Note that this is equivalent to the subset of overall reactions mapped by the RC + 0 rules and is larger than the subset mapped by the Distilled rules. We applied various thresholds to the output atom scores to label them as involved in the catalysis or not. All atoms labeled with the positive class were included in the Learned reaction rule templates.

### Evaluating reaction predictions of reaction rules

To evaluate the reaction rules on their primary task of generating predicted enzymatic reactions, we conducted a test reaction network expansion. The reaction network was seeded with 21 starting compounds designated by the Agile BioFoundry as “beachhead molecules” in a graphic adapted from Lee, S.Y., et al., 2019.^1^ We customized the network expansion tool Pickaxe^12^ to be able to predict reactions featuring multiple unique reactants, and to return atom-mapped reactions. All applicable reaction rules were applied to this starting set of beachhead molecules and then applied again to the union of beachhead molecules and products of the first generation of predicted reactions.

### Calculating evaluation metrics from the test expansion

In addition to simple counts of reactions and compounds, we utilized two metrics to evaluate rules in the test expansion. Reaction feasibility scores were calculated using the DORA-XGB feasibility classifier using its default settings.^46^ We used only the discrete labels and not the scores output by this classifier. Similarity between predicted reactions and known enzymatic reactions from Rhea was calculated as Jaccard similarity on custom extended-connectivity fingerprints.^57^ In addition to the default atom invariants, we included minimal topological distance to any reaction center atom as an atom feature. Atom features are input into an iterative hashing procedure in the usual way. We restricted ourselves to molecular fragments of radius 3 based on all central atoms within 1 bond of the reaction center. In this way, only fragments contained within 4 bonds of the reaction center were represented in the resulting bit vector of length 1024. This choice was motivated by our finding that the vast majority of atoms involved in the catalytic mechanism are found within 4 bonds of the reaction center (Fig. 2).

## ASSOCIATED CONTENT

**Supporting Information**: xxx

## AUTHOR INFORMATION

### Corresponding Author

Linda J. Broadbelt Telephone: +1 847 467 1751

### Author Contributions

S.C.P. – conceptualization of study, design and development of methodology, software development, formal analysis, data curation, investigation, writing (original draft), writing (review & editing), visualization. L.B.J. and K.E.J.T. (equal contributions) – conceptualization of study, provision of computing resources, supervision of research activities, funding acquisition, writing (original draft), writing (review & editing).

## ACKNOWLEDGMENT

This work was performed as part of the Bio-Optimized Technologies to keep Thermoplastics out of Landfills and the Environment (BOTTLE) Consortium and was supported by AMO and BETO under contract no. DE-AC36-08GO28308 with the National Renewable Energy Laboratory (NREL), operated by Alliance for Sustainable Energy, LLC.

Stefan Pate was supported in part by the National Institutes of Health Training Grant (T32GM153505 and T32GM008449) through Northwestern University’s Biotechnology Training Program.

We would like to thank Yash Chainani, Geoffrey Bonnanzio, and Kenna Roberts for helpful discussion.

This research was supported in part through the computational resources and staff contributions provided for the Quest high performance computing facility at Northwestern University which is jointly supported by the Office of the Provost, the Office for Research, and Northwestern University Information Technology.

## DATA AND SOFTWARE AVAILABILITY

All code and data for this work made available at: https://github.com/stefanpate/coarse-grain-rxns

