## Supporting Information for "Mechanism-informed rules tunably balance novelty and feasibility of predicted enzymatic reactions"

Number of pages : 8

Number of figures : 9

Number of tables : 1

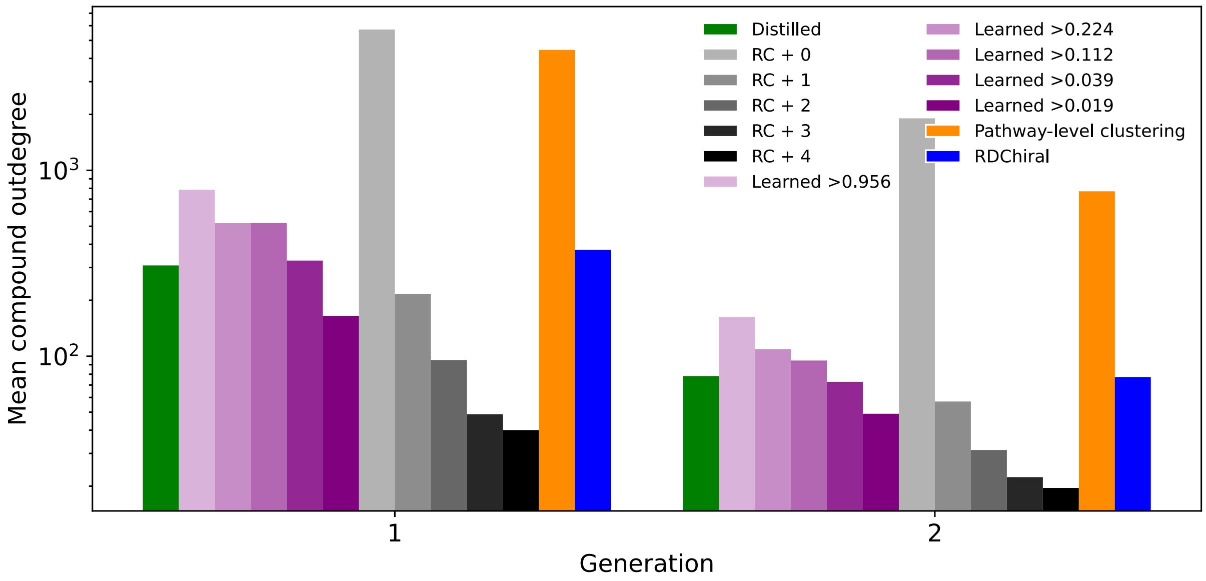


**Figure S1** Average number of reactions predicted using a given compound as a reactant shown for first generation compounds and second generation compounds (products of the first generation reactions).

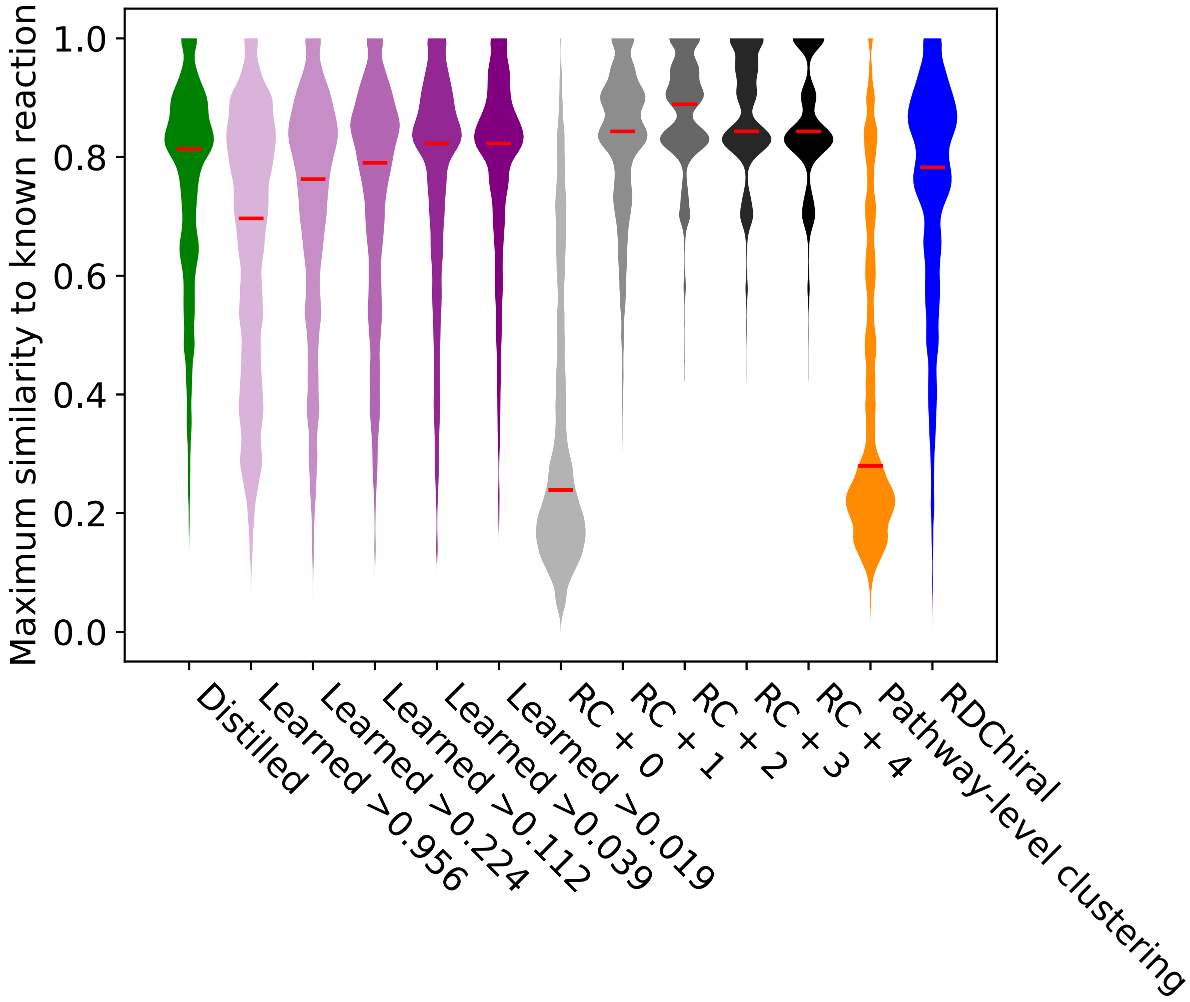


**Figure S2** Distributions of maximum similarity of predicted reactions to known reactions. Similarity is defined as the Jaccard similarity on custom, reaction-center-aware, ECFP bit vectors. See methods for details.

**Table S1** Optimized hyperparameters used in evaluated and production models.

| | **Model #1[1]** | **Model #2[1]** | **Model #3[1]** | **Model #4[1]** | **Model #5[1]** |
|---|---|---|---|---|---|
| **Featurizer mode** | PROD_DIFF[3] | PROD_DIFF[3] | PROD_DIFF[3] | PROD_DIFF[3] | PROD_DIFF[3] |

| | | | | | |
|---|---|---|---|---|---|
| **Hidden vector dimension** | 274 | 247 | 235 | 258 | 293 |
| **# message passings** | 4 | 4 | 4 | 4 | 4 |
| **Prediction head hidden dimensions** | [119, 127] | [156, 191, 117] | [187, 155] | [119, 189, 175] | [138, 167, 196] |
| **Type of prediction head** | FFN[2] | FFN[2] | FFN[2] | FFN[2] | FFN[2] |
| **Batch size** | 103 | 109 | 102 | 105 | 113 |
| **Max epochs** | 6 | 6 | 6 | 6 | 5 |
| **Positive weight scalar** | 0.26 | 0.13 | 0.13 | 0.16 | 0.27 |

[1]Model number corresponds to the outer split of data it was trained on. For production models labeling atoms in overall reactions as an input to Learned rules, we used the average score of the five models.

[2]FFN = feedforward neural network

[3]PROD_DIFF = concatenation of product atom and bond features with the difference of the reactant and product atom and bond features

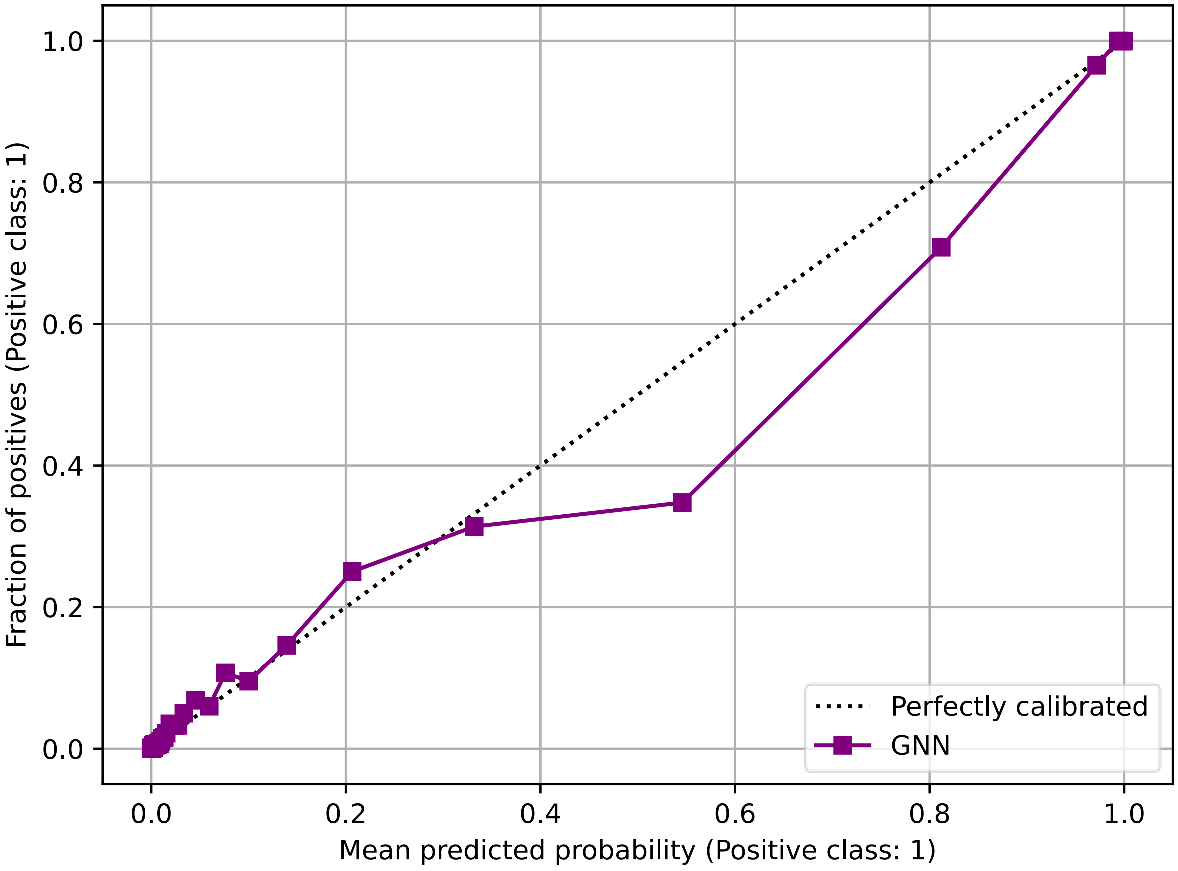


**Figure S3** Calibration curve for GNN classifiers trained on task of labeling atoms required for the catalysis of an overall reaction.

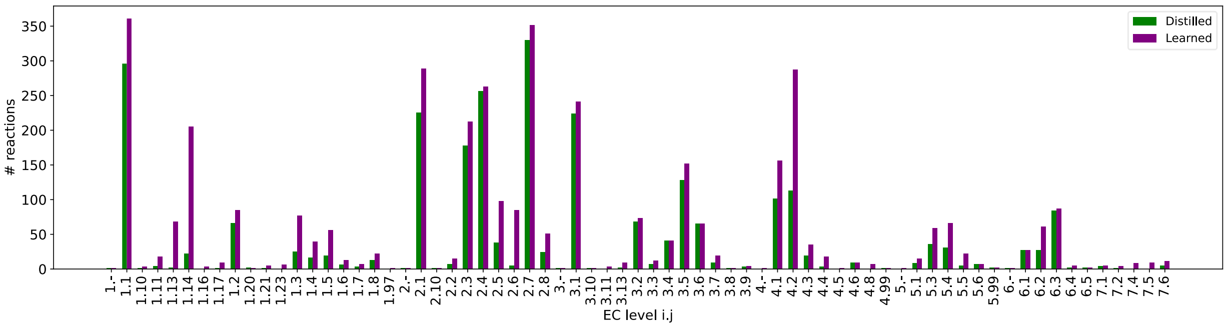


**Figure S4** Coverage of second-digit EC numbers by Distilled and Learned rules.

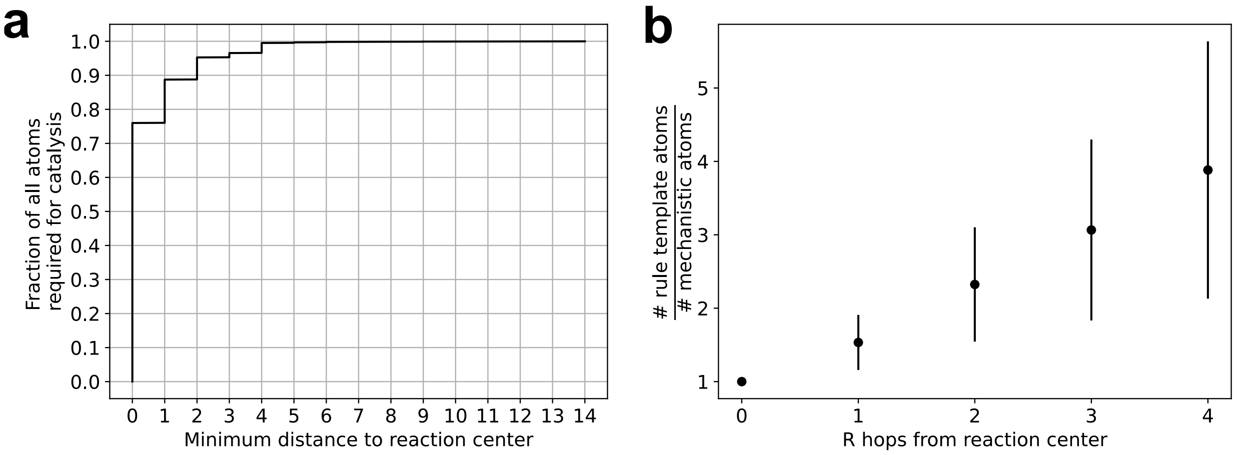


**Figure S5** (a) CDF of fraction of all atoms required for the catalysis of overall reactions in the M-CSA database as a function of minimum topological distance of those atoms to the overall reaction center. (b) Ratio of rule template atoms included in RC + R rules extracted from overall reactions to the number of atoms required for the catalysis of those same reactions as a function of, *R*, the radius around the reaction center included in the templates.

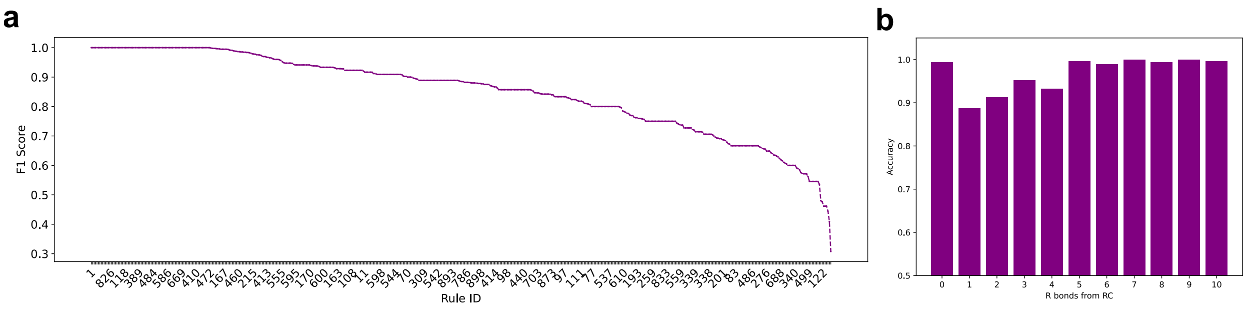


**Figure S6** (a) F1 score on the atom-required-for-catalysis binary classification task grouped by Distilled rule which mapped the reaction. All rules are plotted but only every 10$^{th}$ rule ID is labeled for visualization purposes. Rule IDs uniquely specify Distilled rules in their corresponding CSV file located at the code repository mentioned in the main text. (b) Accuracy against minimum topological distance to the RC, *R*. Both evaluation metrics in (a) and (b) are calculated using an arbitrary decision threshold of 0.5.

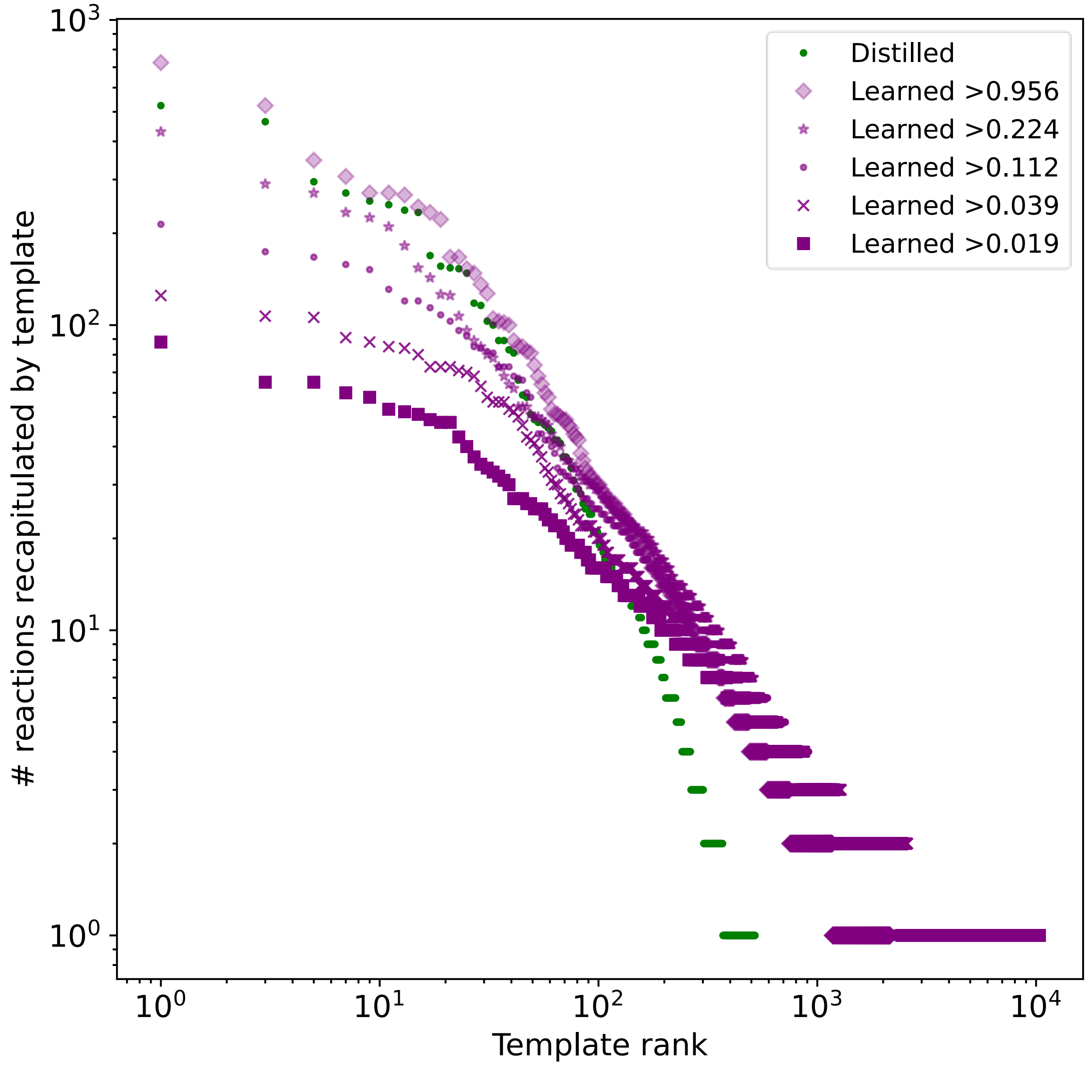


**Figure S7** Distributions of the number of known enzymatic reactions “mapped” (recapitulated) by Distilled and Learned rules. The template rank is simply based on the number of reactions a given template maps, i.e., the template(s) that map the highest number of reactions are ranked 1, and the template(s) that map the lowest number of reactions has the highest rank.

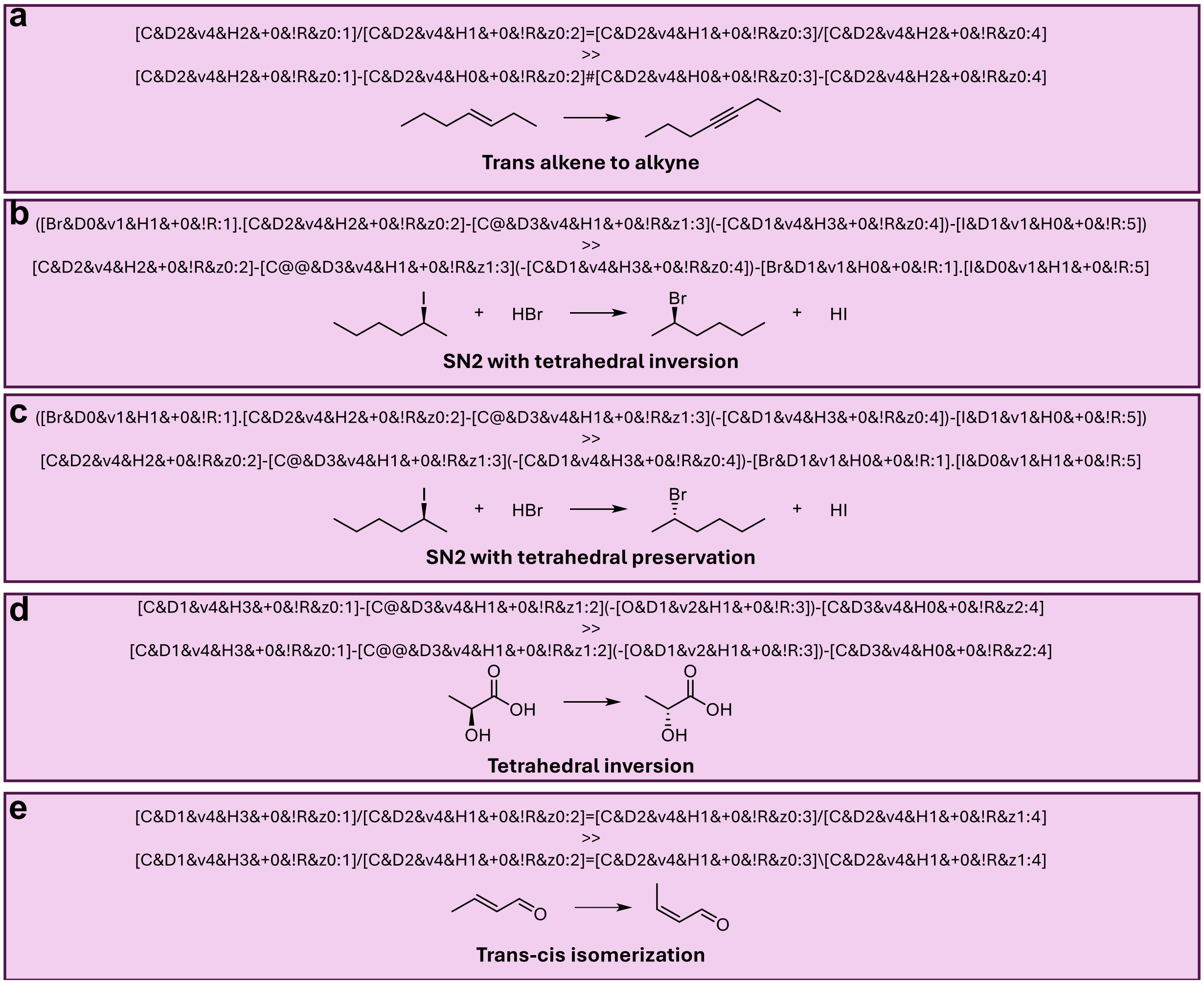


**Figure S8** Template extraction procedure supports stereochemical information. Each box shows an extracted template, reaction, and description. For visualization purposes, the left-hand-side and right-hand-side SMARTS patterns are shown above and below the reaction arrow, respectively. Examples (a)-(c) are taken from Coley et al., 2019. Examples (d) & (e) demonstrate proper application of stereo-only transformations. For all examples, extracted templates are applied to the left-hand-side of the reactions using RDChiral's rdchiralRunText function to ensure the output is equivalent to the right-hand-side of the reaction.

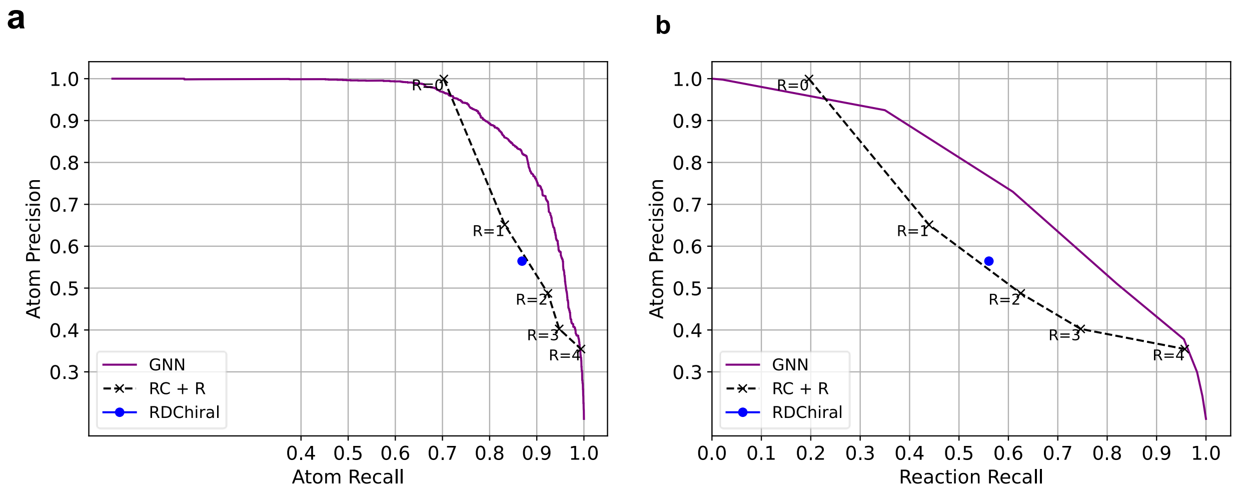


**Figure S9** On reactions exclusively from the M-CSA database, thus with explicitly annotated mechanisms, GNN model successfully labels atoms required for the catalysis of enzymatic reactions. (a) Precision-recall curves for the task of labeling atoms as required for the catalysis of an overall reaction. (b) Precision score evaluated for labeling atoms required for the catalysis versus recall evaluated for full overall reactions. For the latter metric, a correctly recalled data point is defined as an overall reaction with all atoms required for the catalysis of that reaction correctly labeled as with the positive class.
